# Identification of copy number variations and translocations in cancer cells from Hi-C data

**DOI:** 10.1101/179275

**Authors:** Abhijit Chakraborty, Ferhat Ay

## Abstract

**Motivation:** Eukaryotic chromosomes adapt a complex and highly dynamic three-dimensional (3D) structure, which profoundly affects different cellular functions and outcomes including changes in epigenetic landscape and in gene expression. Making the scenario even more complex, cancer cells harbor chromosomal abnormalities (e.g., copy number variations (CNVs) and translocations) altering their genomes both at the sequence level and at the level of 3D organization. High-throughput chromosome conformation capture techniques (e.g., Hi-C), which are originally developed for decoding the 3D structure of the chromatin, provide a great opportunity to simultaneously identify the locations of genomic rearrangements and to investigate the 3D genome organization in cancer cells. Even though Hi-C data has been used for validating known rearrangements, computational methods that can distinguish rearrangement signals from the inherent biases of Hi-C data and from the actual 3D conformation of chromatin, and can precisely detect rearrangement locations *de novo* have been missing.

**Results:** In this work, we characterize how intra and inter-chromosomal Hi-C contacts are distributed for normal and rearranged chromosomes to devise a new set of algorithms **(i)** to identify genomic segments that correspond to CNV regions such as amplifications and deletions (*HiCnv*), **(ii)** to call inter-chromosomal translocations and their boundaries (*HiCtrans*) from Hi-C experiments, and **(iii)** to simulate Hi-C data from genomes with desired rearrangements and abnormalities (*AveSim*) in order to select optimal parameters for and to benchmark the accuracy of our methods. Our results on 10 different cancer cell lines with Hi-C data show that we identify a total number of 105 amplifications and 45 deletions together with 90 translocations, whereas we identify virtually no such events for two karyotypically normal cell lines. Our CNV predictions correlate very well with whole genome sequencing (WGS) data among chromosomes with CNV events for a breast cancer cell line (r=0.89) and capture most of the CNVs we simulate using *Avesim*. For *HiCtrans* predictions, we report evidence from the literature for 30 out of 90 translocations for eight of our cancer cell lines. Furthermore, we show that our tools identify and correctly classify relatively understudied rearrangements such as double minutes (DMs) and homogeneously staining regions (HSRs).

**Conclusions:** Considering the inherent limitations of existing techniques for karyotyping (i.e., missing balanced rearrangements and those near repetitive regions), the accurate identification of CNVs and translocations in a cost-effective and high-throughput setting is still a challenge. Our results show that the set of tools we develop effectively utilize moderately sequenced Hi-C libraries (100-300 million reads) to identify known and *de novo* chromosomal rearrangements/abnormalities in well-established cancer cell lines. With the decrease in required number of cells and the increase in attainable resolution, we believe that our framework will pave the way towards comprehensive mapping of genomic rearrangements in primary cells from cancer patients using Hi-C.

**Availability:**

- CNV calling: https://github.com/ay-lab/HiCnv
- Translocation calling: https://github.com/ay-lab/HiCtrans
- Hi-C simulation: https://github.com/ay-lab/AveSim

## 1 Introduction

Cancer is a disease that is strongly associated with genomic abnormalities and rearrangements, such as CNVs (e.g., amplifications and deletions) and chromosomal translocations (Mitelman et al. 2007; Zack et al. 2013). These often recurrent genomic rearrangements, such as the Philadelphia Chromosome formation in chronic myeloid leukemia (Rowley 1973), are generally associated with certain cancer types and subtypes. Therefore, systematic identification of these rearrangements is critical for understanding the molecular mechanism of oncogenesis as well as for clinical decision making in personalized medicine.

Recent studies have shown that rearrangements in cancer not only alter the one-dimensional (1D) but also the three-dimensional (3D) structure of the genome and individual chromosomes (Engreitz et al. 2012; Ay et al. 2015; Barutcu et al. 2015; Harewood et al. 2017; Seaman et al. 2017). These studies were made possible by the development of high-throughput chromosome conformation capture techniques (e.g., Hi-C) (Lieberman-Aiden et al. 2009; Rao et al. 2014), which also revealed novel relationships between spatial arrangement of the genome and its function (Dixon et al. 2012; Ay et al. 2014; Pope et al. 2014; Sexton and Cavalli 2015). More recent work also link the variation in tightly regulated features of the 3D genome architecture to dysregulation of genes and, increased disease risk in several diseases including cancer (Groschel et al. 2014; Lupianez et al. 2015; Javierre et al. 2016; Lupianez et al. 2016; Schmiedel et al. 2016).

Even though Hi-C data has been used to visually confirm changes in 3D structure due to known genomic rearrangements (Engreitz et al. 2012; Ay et al. 2015; Barutcu et al. 2015; Harewood et al. 2017), to date, there are no computational methods to precisely identify multiple classes of rearrangements *de novo* from Hi-C data. Traditional methods such as PCR, southern blotting and fluorescent in situ hybridization are low throughput, whereas karyotyping is not precise in finding breakpoints of rearrangements (Schrock et al. 1996; Speicher et al. 1996; Davies et al. 2005; LaFramboise 2009). When compared to more recent high-throughput methods for detection of genomic rearrangements, Hi-C has several advantages (Harewood et al. 2017). First, copy number neutral (i.e., balanced) events (e.g., balanced translocations) that are missed by coverage-based methods, such as comparative genomic hybridization (array-CGH) and whole-genome sequencing (WGS), are still detectable by Hi-C. Second, because Hi-C read pairs span all genomic distances (megabases) rather than a fixed insert size, such as 200-800bp, 1kb, 2-5kb for WGS, short-range and long-range mate pair sequencing, respectively, rearrangements involving repetitive regions (e.g., translocations involving a centromere) could be detected by Hi-C from the contact patterns of the surrounding mappable regions. Third, with moderate sequencing depths (150-300M reads or 5-10x coverage), unlike shallow WGS with similar costs, Hi-C allows detection of rearrangements beyond CNVs (Harewood et al. 2017). However, it is important to note that, Hi-C based methods will have limited power to detect fine-scale rearrangements (<50kb in size) due to inherent limitations of the assay (e.g., digestion sites are ∼1-4 kb apart) and/or the cost of sequencing required to achieve sub-10kb resolution contact maps.

Here we provide a set of computational methods, which allow us to predict, simulate and validate genomic rearrangements in cancer with Hi-C data **(Figure 1)**. The inherent challenge in detecting CNVs and translocations from Hi-C data lies in distinguishing the rearrangement related contact patterns from the expected 3D folding of normal chromosomal regions and from technical and experimental biases of the assay. In order to overcome these challenges, here we develop computational methods to detect CNVs (*HiCnv*) and translocations (*HiCtrans*), and, in addition, a versatile pipeline to simulate (*AveSim*) realistic Hi-C contact maps with introduced chromosomal abnormalities. *HiCnv* works on contact counts at the single restriction enzyme (RE) fragment level in order to leverage Hi-C data at its highest possible and native resolution. Briefly, *HiCnv* first computes 1D read coverage for each RE fragment, followed by normalization, smoothing (kernel density estimation, KDE) and segmentation (Hidden Markov Model, HMM). The CNV segments are further processed for refinement of their breakpoint coordinates (segment ends) and assignment of their CNV labels **(Figure 1).** *HiCtrans*, on the other hand, scans fixed-size resolution (e.g., 40kb). interchromosomal contact maps of each chromosome pair for potential translocations using change-point statistics **(Figure 1).**

**Fig. 1.**
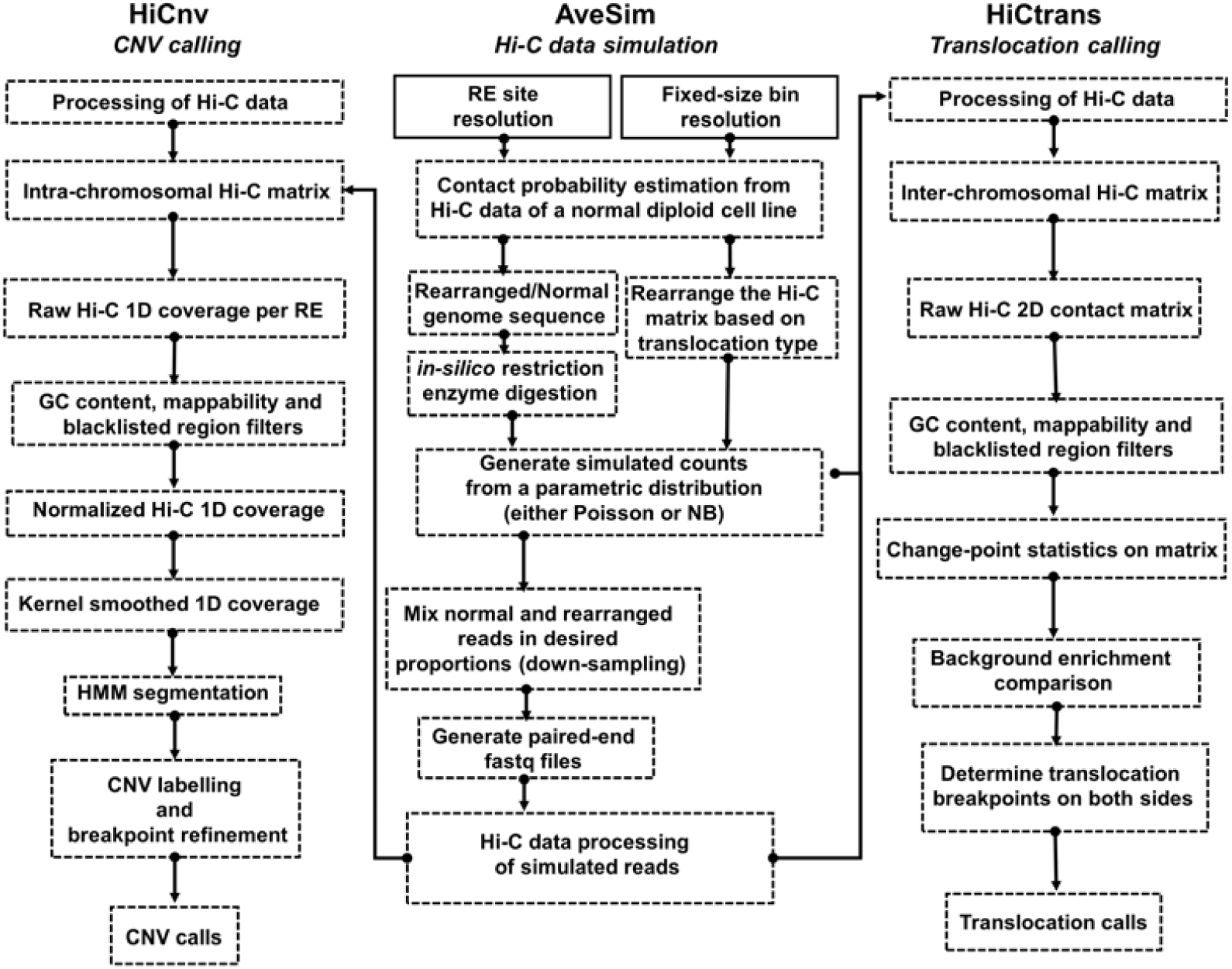
Overall summary of the methods developed in this work. *HiCnv* and *HiCtrans* are the two methods we develop for identifying copy number variations (CNV) and translocations from Hi-C data. *AveSim* is our simulation pipeline that generates simulated Hi-C data with genomic rearrangements introduced. *AveSim* works either at restriction enzyme (RE) site/fragment resolution or with fixed-size genomic bins to create intra- and inter-chromosomal Hi-C contact maps with the desired rearrangements in the genome.

Our results suggest that *HiCnv* and *HiCtrans* can accurately predict large scale CNVs (>1Mb) and translocations from both real Hi-C data of cancer cell lines and in simulated contact maps with desired rearrangements. We compare our CNV calls with existing sequencing-based (WGS) and array-based (HAIB) genotyping efforts and show that our results largely agree with WGS and to a less r extent with HAIB. The agreement between WGS and HAIB CNV calls were also low. Both HiCnv and WGS results are consistent with previous karyotyping at the chromosomal level. We show that our CNV segment breakpoint predictions are at a median distance of 3 RE sites (∼12kb) from simulated breakpoints and are enriched for transition of WGS coverage scores. For translocations, we identify a total of 90 translocating chromosome pairs in our 10 cancer cell lines giving rise to 227 distinct contact enrichments in inter-chromosomal maps. We find evidence in the literature for 30 of our 90 reported translocations. Finally, we report two highly amplified regions that we predict as double minutes through simulations of expected contact patterns. Our results demonstrate the power of Hi-C data in predicting a wide range of large scale genomic rearrangements and the importance of developing flexible tools to simulate Hi-C data with custom made genomic rearrangements.

## 2 Methods

### 2.1. Data Collection

We download raw reads for publicly available Hi-C data for 12 cancer cell lines from the ENCODE project portal (encodeproject.org) as well as from two normal diploid cell lines, namely hESC (Dixon et al. 2012) and IMR90 (Jin et al. 2013). We discard two cancer cell lines due to concerns about Hi-C data quality and mapping rates (SJCRH30 and SKNDZ) **(Table 1)**. All of these Hi-C libraries were generated using the 6-bp cutter HindIII restriction enzyme (cut site: A|AGCTT). We process the raw reads for each cell line with HiCPro (Servant et al. 2015) to generate raw intra- and inter-chromosomal matrices at a single RE resolution and with fixed-size bins (e.g., 40kb). For comparing our CNV calls, we use a ∼30x coverage WGS data processed at 50kb bins for the T47D cell line (personal communication with Feng Yue). We also use CNV calls from ENCODE Illumina BeadChip array data generated by Hudson Alpha Institute (HAIB) and processed by circular binary segmentation (CBS) (Olshen et al. 2004) for four of our cancer cell lines, namely T47D, PANC1, LNCaP and A549 (wgEncodeHaibGenotype). For known translocations, we use karyotype information from several websites as well as previous publications about each cell line. We perform all of our analysis using the human genome build GRCh37 (hg19).

**Table 1.**
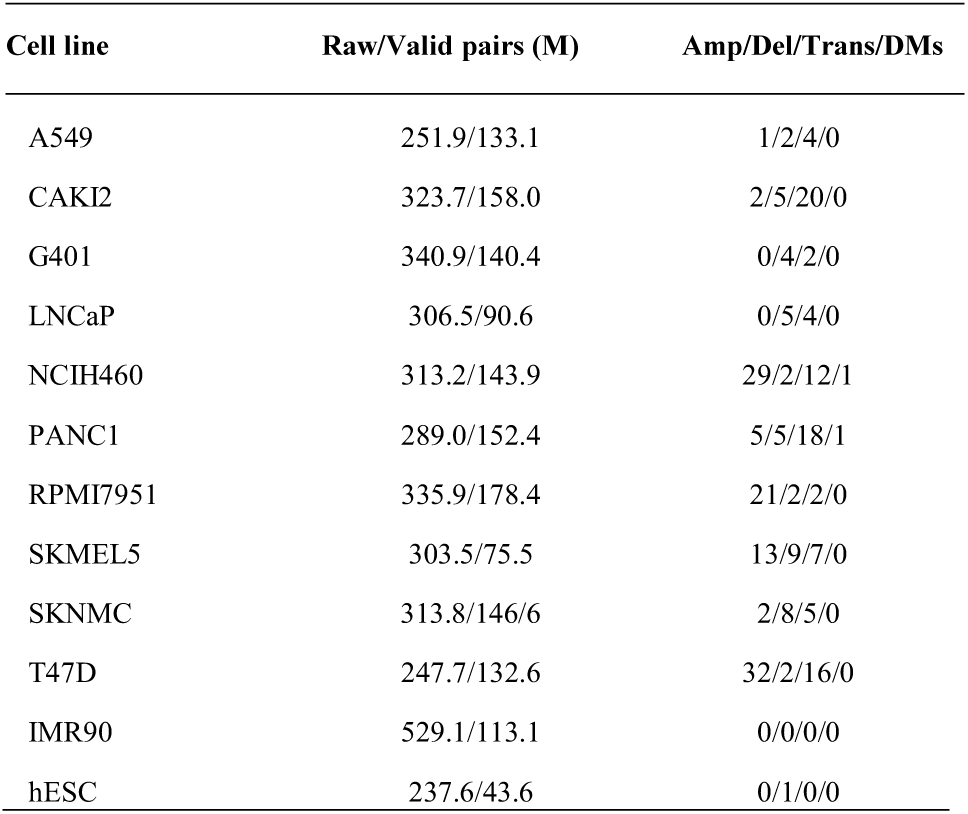
Summary of Hi-C data used in this work. The number of reads, valid pairs (in millions); amplifications (Amp), deletion (Del), translocations (Trans) and double minutes (DMs) are given.

### 2.2. Copy Number Variation calling from Hi-C (*HiCnv*)

For calling CNVs, we first compute Hi-C read coverage at each restriction site throughout the genome. Since all of our samples are digested using HindIII, we get coverage measurements on average from every 4kb of the genome. This restriction site based approach, instead of fixed size genomic bins allows us to utilize Hi-C data at its native and highest possible resolution. We filter out RE fragments with low GC-content (<0.2), mappability (<0.5) or any overlap with blacklisted genomic regions as prescribed by ENCODE (Consortium 2012). Since the CNV information is reflected in a region’s coverage, commonly used Hi-C normalization methods that aim to equalize coverage (Imakaev et al. 2012; Rao et al. 2014) are not appropriate because they cancel out the copy number aberrations **(Supp. Fig. 1)**. Therefore, starting from a previously described regression-based method, HiCNorm (Hu et al. 2012), here we employ a one-dimensional regression-based normalization (1Dreg) on the RE fragment level coverage counts that corrects for factors that are known to bias Hi-C data (i.e., GC content, mappability, fragment length) without removing the CNV signal (**Figure 2**). Since it works on 1D coverages rather than 2D contact maps, our regression is scalable to RE fragment level data. For a given chromosome, let *c*_*i*_ denote total number of read ends that map to the *i*^th^ RE fragment where *i=1..n* and *frag*_*i*_*, gc*_*i*_ *and map*_*i*_ represent the effective fragment length, GC content and mappability of the fragment *i.* The equation for our Poisson regression model is as follows:

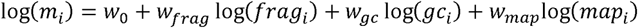

**Fig. 2.**
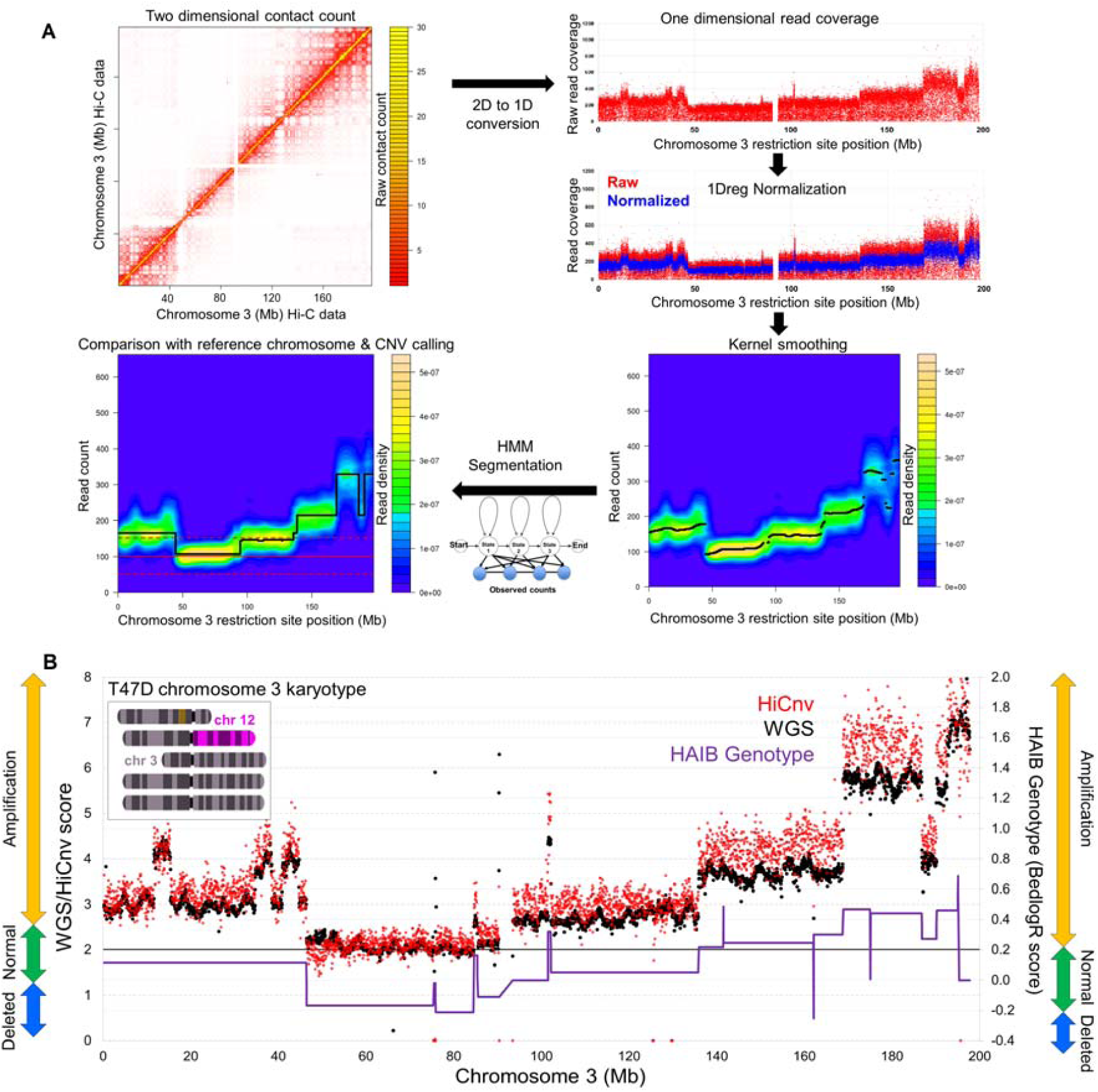
Identification of CNVs from Hi-C data. A) Our *HiCnv* pipeline starts with extracting a 1D coverage vector for each RE site from the intra-chromosomal contact matrix. We then use a 1D Poisson regression model (1Dreg) to normalize the raw coverages for GC content, mappability and fragment length biases, and apply kernel smoothing to further remove noise. Next, we use an HMM-based approach to find the possible CNV segments from the normalized and smoothed coverage values. We assign CNV labels such as amplified or deleted by comparing the mean segment coverage against that of a diploid reference chromosome. We also correct for edge effects due to smoothing to refine our CNV breakpoint calls. **B)** Comparison of HiCnv (red) and WGS (black) coverage scor s for chromosome 3 of T47D (Pearson corr. 0.92). We also plot the bedlogR scores and CNV labels for the same chromosome obtained from HAIB genotype (WGS vs HAIB bedlogR corr. 0.81). HAIB CNV labels largely disagree with both *HiCnv* and WGS, whereas HiCnv and WGS labels agree with each other and with karyotyping (top left inset).

Here *w*_*0*_ denotes the intercept, *w*_*frag*_*, w*_*gc*_ and *w*_*map*_ represent the fragment length, GC content, and mappability biases, respectively. We assume that *c*_*i*_ follows a Poisson distribution with rate *m*_*i*_ and estimate the normalized count for RE fragment *i* as the residual *r*_*i*_*= c*_*i*_ /*m*_*i*_. **Supp. Fig. 2** shows that 1Dreg effectively removes aforementioned biases but not the differences between regions with different copy numbers.

Next, we use the normalized coverage values for bivariate kernel density estimation, which provides a probabilistic measure to find a certain magnitude of count for each RE site in a given chromosome. We use bkde2d function in R (Wand 1994) for KDE as follows:

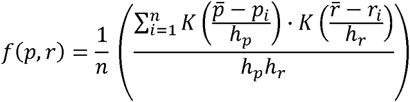

Here p_i_ and *r*_*i*_ denote the index and the normalized coverage of an RE site, where *i=1..n. K* is the standard Gaussian probability density function, and *h*_*p*_, *h*_*r*_ correspond to x-axis and y-axis bandwidths used for KDE, respectively. *f(p, r)* gives the final probability density of observing contact counts surrounding an RE position in the genome. We then assign each RE site the count that is associated with highest density in the density estimate. **Supp. Fig. 3** shows that KDE is an essential step as it eliminates much of the unwanted noise from the Hi-C reads prior to the segmentation process. Since the choice of bandwidth parameter for the x-axis (number of consecutive RE sites) has a significant effect on the number of false positive and false negative identifications, we assess different band-width values on simulated data **(Supp. Fig. 4)**.

Consequently, we use a Hidden Markov Model-based approach (HMM) to perform a segmentation on the KDE smoothed RE site coverage values. We use the RHmm package (https://github.com/cran/RHmm) to fit a univariate-Gaussian model to our coverages in order to identify change points in the data. We use Bayesian information criterion (BIC) to select the optimum number of states in input coverages and subsequently the mostly likely hidden state that correspond to copy number variation status. However, since cancer cell lines can have very high modal numbers (e.g., hyper-triploid HeLA cell line), assigning CNV labels such as deleted/normal/amplified is not trivial because the genome-wide average may not reflect a truly diploid reference point. We, thus, choose a reference chromosome based on available biological information such as whole genome sequencing or karyotyping information as a baseline to determine the true copy number of each chromosome and accurate labelling of the segmented regions within a chromosome. Even though finding such a reference chromosome is possible for all of our cell lines and can be partially automated, it should be noted that, only our whole chromosome level estimates are dependent on it (i.e., distinguishing two full copies of a chromosome from three full copies). Our segmentation within each chromosome can label CNVs as amplified or deleted relative to each other without a reference.

Next, we assign CNV labels by calculating an average normalized coverage per RE site for each CNV segment *i*, *NC*_*i*_. We then divide this number by the sum of all such coverages across all chromosomes to get a proportion of interaction count, *PIC*_*i*_, per chromosome or per a given segment. In the case of a given diploid reference chromosome *ref*, we determine CNV labels by comparing the segment mean *PIC* value to *PIC*_*ref*_. We use a percentage threshold on the increase or decrease of a segment’s mean with respect to the reference in order to label it as an amplification or deletion. We assess different threshold values in **Supp. Fig. 4.** Without a reference chromosome, the same process could be followed using the average *PIC* value either across all chromosomes or within each chromosome.

One important step remaining after labeling CNVs is the refinement of segment breakpoint coordinates, which may be off because of edge effects introduced by the KDE smoothing **(Supp. Fig. 5)**. Such effects are known to be present at the segment boundaries, where the coverage is discontinuous and in transitive phase from one segment to the next (Chiu 2000). In order to detect precise breakpoint coordinates of a CNV segment, we compute the ratio between the median coverage of 10 upstream and 10 downstream RE sites for each individual RE within a distance less than the selected bandwidth from each segment end. We report RE sites with peaks or dips of this ratio as the breakpoints between two adjacent segments with differing CNV labels **(Supp. Fig. 5)**.

### 2.3. Translocation calling from Hi-C data (*HiCtrans*)

In order to detect translocation events, we analyze the inter-chromosomal contact matrices at a fixed-size bin resolution (40kb in this work) for each possible pair of chromosomes from a given cell line. For a given pair of chromosomes, we perform binary segmentation first on each row and then on each column of the inter-chromosomal Hi-C matrix of that chromosome pair **(Figure 3)**. For binary segmentation, we use the *BINGSEG* function from the *changepoint* package in R, which runs very fast and reports more than one single segment only when a significant change is detected in a given contact count row/column (Scott 1974). This allows us to efficiently scan the full inter-chromosomal contact matrix to detect candidate change-point events from the perspective of each chromosome.

**Fig. 3.**
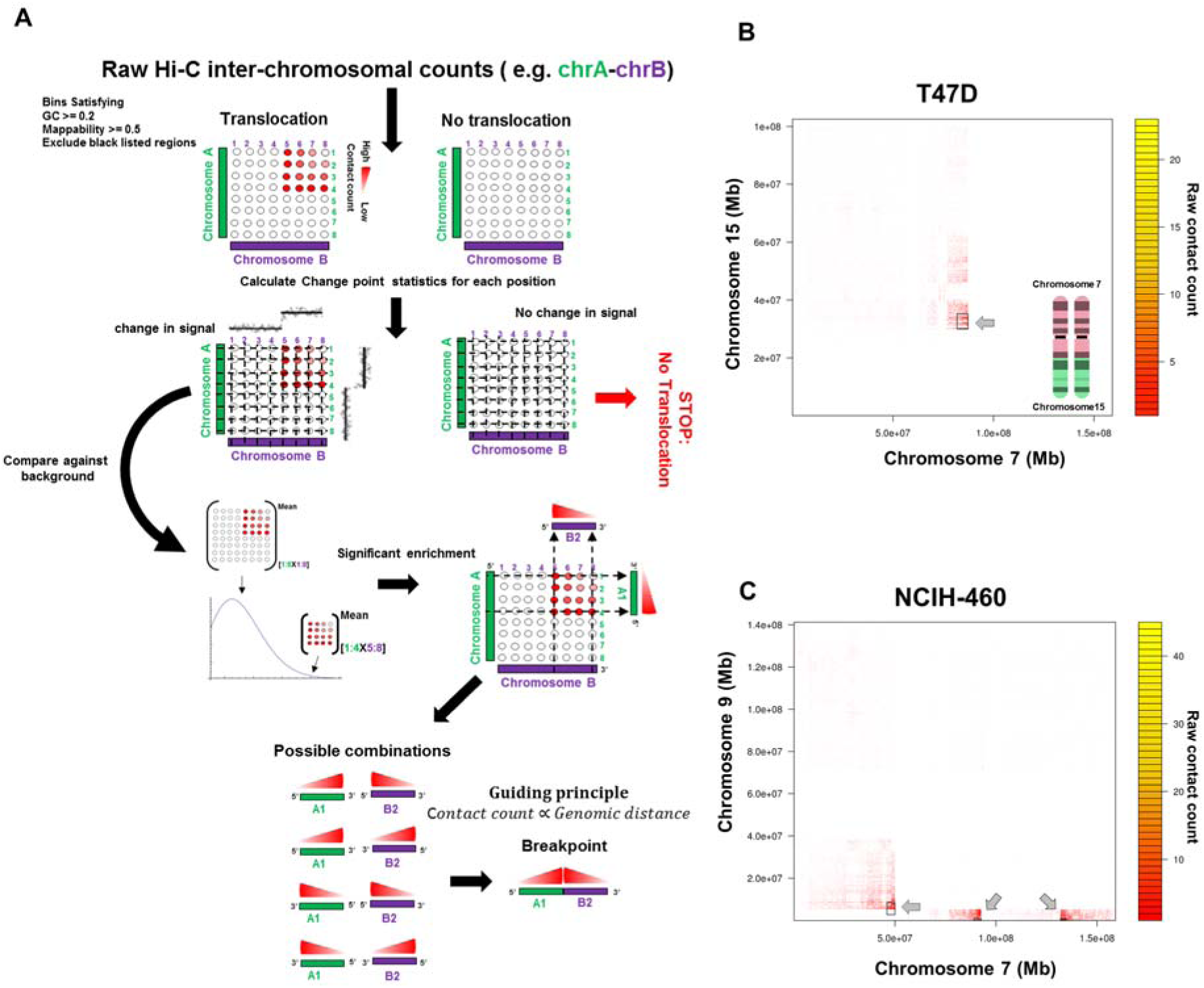
Identification of translocations from Hi-C data. A) Our *HiCtrans* pipeline starts with performing binary segmentation independently on each row and column of an inter-chromosomal matrix. *HiCtrans* then aggregates the change points identified from the perspective of each chromosome to determine rectangular boxes of contact enrichment with respect to the overall inter-chromosomal matrix. For each box, *HiCtrans* computes a statistical significance for enrichment and for boxes passing this significance test, it reports the pair of regions with the highest contact count as the breakpoints. Two previously validated/observed real case examples of translocations identified by *HiCtrans* in **B)** T47D and **C)** NCIH460 (lung cancer) cell line.

Next, we sequentially scan each row for the existence of a change point. Let row *i* be the first such row with a change point at coordinate *j.* We then determine an interval of [*i, i+x-1*], which consists of consecutive rows that all have a change point at *j.* We perform a similar operation on the columns of the interchromosomal matrix, which provides us another interval [*j, j+y-1*] of consecutive columns with a common change point from the interval [*i, i+x-1*]. This provides us with a rectangular box of size *x* by *y* in the interchromosomal matrix spanning *x* rows of the first chromosome and *y* rows of the second chromosome that are in consideration. These boxes represent regions of inter-chromosomal contact enrichment, which may correspond to translocation events. We then test the significance of this enrichment within a given box compared to the overall inter-chromosomal matrix by comparing the mean contact counts of the two using a Poisson test. We then correct the resulting p-value for the number of such boxes tested for a given chromosome pair and consider further only boxes with an adjusted p-value less than 0.05. This ensures filtering of spurious boxes from the segmentation that do not have significant enrichment of contacts expected as a result of a real translocation event. For each box that passes the significance test, we then find the maximum inter-chromosomal count within that box and report the corresponding row and column as the breakpoints at which the two translocating regions ligate to each other. We further eliminate weak inter-chromosomal signals, by only reporting boxes with a maximum count greater than five as our translocation breakpoints.

### 2.4. A versatile pipeline for simulating Hi-C data with genomic rearrangements (*AveSim*)

In order to provide a benchmark for our HiCnv and HiCtrans calls, we develop a simulation pipeline for generating Hi-C data from chromosomes with inserted genomic rearrangements. We carry out these simulations either at the RE site resolution or with fixed-size genomic bins, which are described in detail below.

#### Simulating rearrangements at the RE fragment level

For the amplification, deletion and translocation simulations, we first generate a rearranged sequence with an extra copy, deleted or translocated region of a chromosome, respectively **(Figure 1, Supp. Fig. 6-9)**. The rearranged sequence is then *in silico* digested with desired restriction enzyme (e.g., HindIII) to get the all possible RE sites. We use Hi-C data from a karyotypically normal cell line (e.g., IMR90) to learn scaling of contact probability with respect to genomic distance using a smooth spline fit. We then use the selected probability distribution to draw random contact counts among the pairs of RE sites in the rearranged genome using their genomic distance, and generate paired-end reads from 50bp regions randomly selected within 500bp of each RE site. This pipeline also allows us to mix and match user defined proportions of reads from the normal and rearranged genomes in order to simulate homogeneous (e.g., 10M reads from the rearranged copy) and heterogeneous rearrangements (e.g., 5M reads each from the normal and rearranged copies, 50% rearrangement) **(Supp. Fig. 6-9)**.

#### Simulating translocations with fixed-size genomic bins

Apart from simulating at RE site resolution, we also carry out translocation simulations with fixed-size genomic bins. For this, we generate different biologically relevant translocation types, namely reciprocal-balanced, non-reciprocal-balanced and unbalanced **(Figure 1, Supp. Fig. 10-11)**. For each type, we first choose a pair of chromosomes and a pair of genomic positions to be exchanged/translocated. Next, we simulate intra- and inter-chromosomal Hi-C contact counts based on the assumption that exchanged genomic regions will interact with their new neighboring regions in a distance-dependent manner similar to real intra-chromosomal neighbors and that they will interact with their previous locations similar to what is expected from inter-chromosomal partners. In order to generate these simulated counts, we use contact probabilities from either Poisson or negative binomial (NB) distributions by determining the best fit using Bayesian information criterion (BIC). We use NB for intra-chromosomal counts as it is the better fit for distances up to 100Mb, and Poisson for inter-chromosomal counts **(Supp. Fig. 10-11).**

## 3 Results

### 3.1. Evaluation of CNV predictions from *HiCnv*

We first ask how *HiCnv* prediction performance is affected by selection of the KDE bandwidth and CNV labeling threshold parameters. For this, we generate a total of 508 amplifications and 453 deletions with sizes ranging from 1Mb to 30Mb in randomly selected regions genome-wide **(Supp. Fig. 4)**. We observe that the optimal KDE bandwidth value estimated by the *bkde2* function outperformed two other bandwidth values in terms of both low false positive and false negative rates **(Supp. Fig. 4).** Among CNV labeling thresholds, we observe that 40% and 45% give balanced results compared to smaller percentages with higher false positive rates and compared to 50%, which has a very low recall. With these selected parameter settings (optimal bandwidth and 45% threshold) and a stringent criterion for defining true positives (predicted CNV needs to cover 100% of the simulated), we correctly identify 313 and 291 simulated amplifications and deletions with only 3 and 47 false positives, respectively(**Supp. Fig. 4)**. For the correctly identified CNV segments, we predict segment boundaries very accurately with a median offset of 3 RE sites (∼12kb). Since the simulated CNVs were scattered across the genome, we get varying performance depending on the genomic features and coverage variation of the region of CNV insertion.

We then run HiCnv on 10 cancer and 2 normal cell lines resulting in a total of 105 amplifications and 45 deletion calls (**Table 1**, **Supp. File 1**). The only CNV reported from the normal cell lines was the whole X chromosome as a deletion in hESC, which is expected as it was derived from a male donor. We then compare our calls with WGS data from T47D, which showed that HiCnv coverages and CNV calls highly agree with the WGS scores at the genome-wide level (**Supp. Fig. 12**), at the level of whole chromosomes (**Figure 4A)** and CNVs within individual chromosomes (**Figure 2B, Figure 4B**). Across all chromosomes with CNVs, HiCnv and WGS scores show 0.89 overall correlation and an average correlation of 0.83 with a standard deviation of 0.11 per chromosome **(Supp. Fig. 12)**. Similar to simulation results, we also assess whether our segment breakpoint calls are accurate in T47D by making aggregate plots of WGS scores around the coordinates of HiCnv predicted breakpoints between normal and amplified regions. Our results show that the refinement step of HiCnv successfully corrects edge effects and reports breakpoints that are at the transition of WGS scores **(Figure 4C)**.

**Fig. 4.**
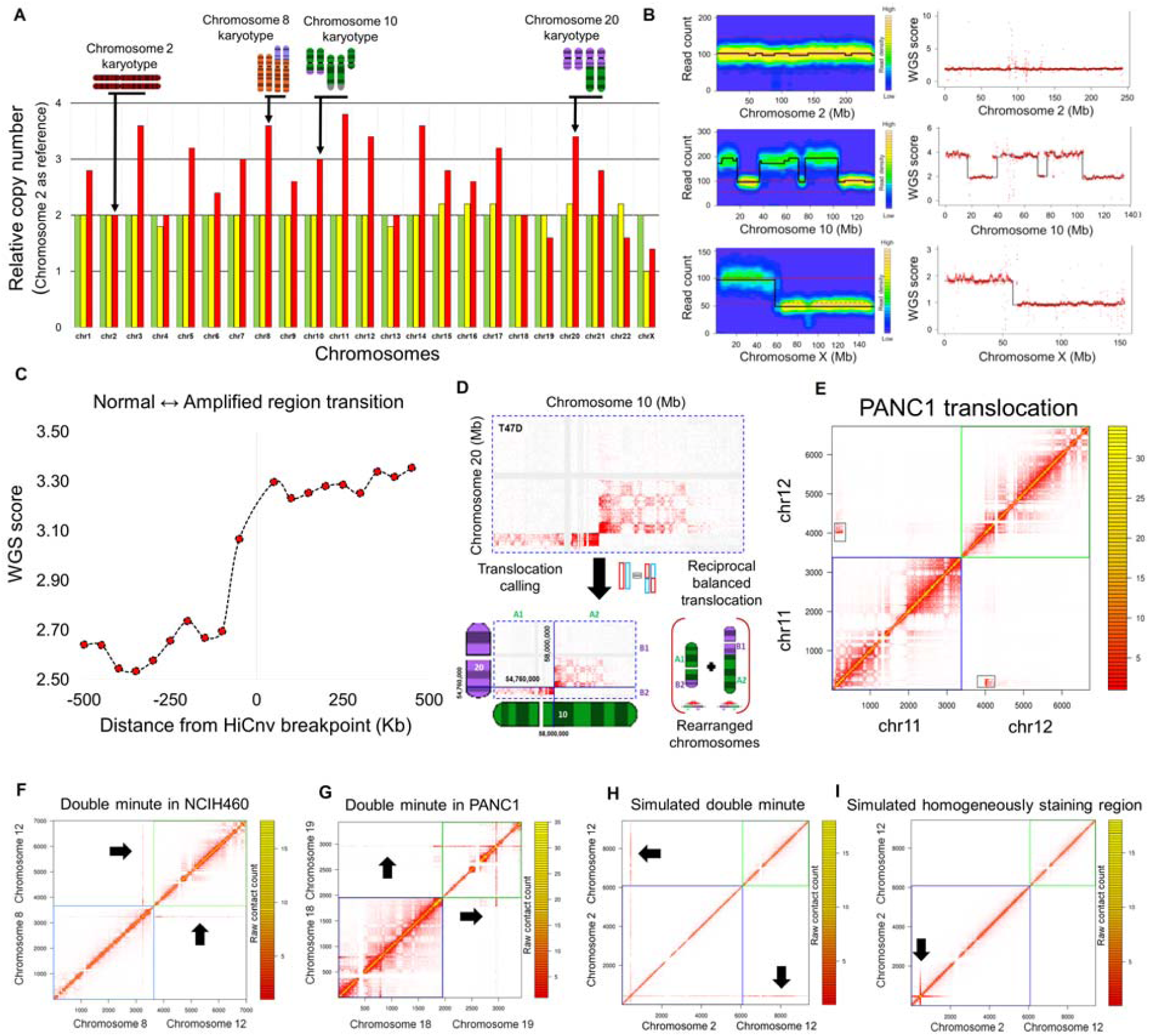
Summary of results. A) Relative copy numbers of each human chromosome in one cancer (T47D) and two karyotypically normal (IMR90: fibroblast, hESC: embryonic stem cells) cell lines computed by HiCnv using chromosome 2 as our diploid reference chromosome. Karyotype information from T47D is shown for a subset of chromosomes. For chromosomes with CNVs other that whole chromosome deletions or amplifications in T47D, the relative copy number reflects chromosomal average (e.g., chr1, chr3, chr10, chr20) rather than true copy numbers (e.g., chr2, chr4, chr5, chr13). Both IMR90 and hESC relative copy numbers are 2+/-0.2 confirming that all chromosomes have two copies except for chromosome X in hESC, which was derived from a male donor. **B)** Read coverage plots for three example chromosomes and their corresponding segmentations using *HiCnv* from T47D Hi-C data (left panel) and T47D WGS data (right panel). **C)** The distribution of T47D WGS scores flanking the HiCnv identified breakpoints between adjacent CNVs. We orient 47 such transitions such that all transition go from a normal to an amplified region and plot the median value of each 50kb region within +/-500kb of the identified breakpoint. **D)** An example inter-chromosomal Hi-C contact map from the breast cancer cell line T47D with a known translocation (Davidson et al. 2000) between chromosomes 10 and 20, and a depiction of reported breakpoint coordinates and rearrangement reported by *HiCtrans* (left). **E)** Heatmap of a known translocation (Espino et al. 2009) in a pancreas cancer cell line (PANC1) identified by *HiCtrans.* **F-G)** Heatmaps depicting the contact patterns generated by the two highly amplified regions we identify in NCIH460 and PANC1 cell lines that are of size 1.71Mb (chr8) and 1.07Mb (chr19), respectively. **H-I)** Two panels show contact patterns generated by AveSim for an introduced double minute (DM) and a homogeneously staining region (HSR), respectively, suggesting that both of the highly amplified regions in **F** and **G** are DMs and not HSRs.

We also use compare our CNV calls to those from HAIB genotype for four cancer cell lines **(Figure 2B, Supp. Fig. 13)**. Our results show that the overlap between segments from the two sets highly varies among cell lines and depends heavily on whether its coverage is computed from HiCnv or HAIB segments. For instance, for T47D cell line, 30 of the 31 regions identified as amplified by HAIB are fully contained in HiCnv calls, whereas only 7 out of the 26 HiCnv amplifications were covered by HAIB. **Figure 2B** provides a potential explanation to the low number in the latter case as it shows that HAIB CNV labels can be incorrect due to overestimation of “normal” for chromosomes that are mainly amplified. For the example in **Figure 2B,** CNV labels from HAIB also disagree with WGS and karyotyping results, even though the HAIB scores correlate well with WGS (r=0.81).

### 3.2. Evaluation of translocation calls from *HiCtrans*

A translocation event introduces one or more regions of contact enrichment depending on the type of the translocation **(Supp. Fig. 9, 11)**. These enrichment regions correspond to breakpoints, at which two previously inter-chromosomal regions ligate to one anoth r **(Supp. Fig. 8, 10).** Here we capitalize on two observations: 1) due to formation of chromosome territories, only a very small fraction of interactions occur between chromosomes compared to within, 2) for a pair of chromosomes with a translocation, the inter-chromosomal Hi-C matrix exhibits contact enrichments that are comparable to intra-chromosomal contacts and are distance dependent. Our method HiCtrans identifies and reports these contact enrichments (i.e., breakpoints) as the inter-chromosomal translocation signatures **(Figure 3A)**. To evaluate predictions by HiCtrans, we first simulate different types of naturally occurring translocations through our AveSim pipeline. We show that HiCtrans accurately identifies the exact breakpoint locations of 176 translocations with 12 false positives from a total of 200 simulations of reciprocal and non-reciprocal balanced translocations carried out at 40kb resolution **(Supp. Fig. 10).**

Next, we assess the robustness of our translocations calls to noise or heterogeneity in Hi-C data. For this purpose, we use *Avesim* to generate RE site level simulations, which create paired-end read libraries that contain a mixture of translocated and normal copies of the chromosomes **(Supp. Fig. 6-9)**. Our results in this setting show that HiCtrans is very robust to heterogeneity and can accurately identify translocations even when only 5% of the reads come from translocated copies **(Supp. Fig. 9B).** This suggests HiCtrans could detect low prevalence sub clonal translocations within, even though we do not know how the success at 5% from our simulations would translate into real sub clonal events.

We then apply HiCtrans to our 10 cancer cell lines, which resulted in identifications of 90 translocations corresponding to a total 227 contact enrichments **(Table 1, Supp. File 2)**. Even though there are no comprehensive sets of validated translocation calls, previous studies on A549 (lung), LNCaP (prostate), NCIH-460 (lung), PANC1 (pancreas), SKMEL5 (skin), SKNMC (brain) and, T47D (breast) cancer cell lines identified several translocations, against which we compare our calls (Takahashi et al. 1989; Davidson et al. 2000; Espino et al. 2009; Peng et al. 2010; Rondon-Lagos et al. 2014). We find that 30 out of our 90 translocations were previously reported **(Supp. File 2)**. One of these examples is a non-reciprocal translocation between chromosomes 7 and 15 in T47D cells (Davidson et al. 2000), which results in two copies of t(7:15) translocation products, two copies of normal chromosome 7 and one copy of the normal chromosome 15 **(Figure 3B)**. Another example is the formation of t(7;9) (Takahashi et al. 1989), which is a marker chromosome of the NCIH-460 non-small cell lung cancer cell line **(Figure 3C)**. **Figure 4D-E** show two additional translocations that are also characterized before (Davidson et al. 2000; Espino et al. 2009). We also show contact patterns from a number of other known translocations in **Supp. Fig. 14**, and in **Supp. Fig. 15** we provide similar heatmaps from 12 translocations that are novel predictions by HiCtrans to the best of our knowledge. Even though further experiments (e.g., FISH, PCR) are needed to confirm these potentially novel translocations, we believe that the similarity of contact patterns of known translocations to these new predictions provides strong evidence for their presence in these cell lines.

## Discussion

Here we proposed two new methodologies (*HiCnv* and *HiCtrans*) that utilize chromatin contact maps to identify CNVs and translocations in cancer cell lines. Our comparisons with whole genome sequencing, HAIB genotype and karyotyping demonstrated that our methods accurately detect large scale CNVs (>1Mb) and translocations from moderately sequenced Hi-C libraries, thereby, extending the utility of Hi-C data to detection of genomic rearrangements in cancer.

Due to lack of precise maps of genomic rearrangements, which could be used as true gold standards for evaluating our predictions, we also developed a simulation framework (*AveSim*) that can introduce rearrangements in Hi-C contact maps. This framework provided us with two different approaches to generate simulated Hi-C data as well as the ability to introduce several different types of naturally occurring rearrangements into contact matrices. An interesting example was the highly amplified regions identified in two cancer cell lines NCIH460 (chr8:128691158-130402069) and PANC1 (chr19:39774739-40847406) **(Figure 4F-G)**. Such aberrant amplifications have been reported previously in neuroblastoma (Balaban-Malenbaum and Gilbert 1977; Storlazzi et al. 2010) and fall into two classes depending on whether the amplified regions are localized to their native locus (homogenously staining regions –HSRs) or created extrachromosomal DNA loci (double minutes –DMs). By simulating both of these possible scenarios **(Figure 4H-I)**, we show that both regions we identify produce contact patterns consistent with DMs and not with HSRs. This example highlights the power of Hi-C data and the importance of a versatile simulation method, in not only evaluating the performance and tuning the parameters of developed methods, but also in generating signature contact patterns from complex rearrangements. Such complex rearrangements remain difficult to identify with any high-throughput method including Hi-C.

We believe that the framework we put forward here for identifying genomic rearrangements solely from Hi-C data, as well as a pipeline for simulating such rearrangements in Hi-C contact maps, which can be used as a benchmark for predictions methods developed here and later on, is valuable to the fields of cancer research and 3D/4D nucleome. We believe one future direction is to develop probabilistic methods that will utilize higher resolution Hi-C datasets to identify CNVs that only occur in a subset of cells from a potentially heterogeneous tumor sample. Another important direction is the integration and reconciliation of rearrangement calls from multiple high-throughput experiments, such as conventional whole genome sequencing (WGS), linked-read WGS, single-molecule imaging (Bionano Irys) and Hi-C (Zheng et al. 2016; Dixon 2017), in order to come up with comprehensive and accurate maps of genomic rearrangements in cancer.

## Acknowledgements

The authors would like to thank ENCODE 3D Nucleome Working Subgroup members David Gilbert, Job Dekker, William S. Noble, Feng Yue, Jesse Dixon and members of their groups for valuable discussions and Feng Yue’s group for sharing their unpublished whole genome sequencing data for the T47D cell line.

